# A workflow for simultaneous detection of coding and non-coding transcripts by ribosomal RNA-depleted RNA-Seq

**DOI:** 10.1101/2021.01.04.425201

**Authors:** Nikita Potemkin, Sophie M.F. Cawood, Jackson Treece, Diane Guévremont, Christy J. Rand, Catriona McLean, Jo-Ann L. Stanton, Joanna M. Williams

**Author notes:** Correspondence to: Assoc. Prof. Joanna Williams, Department of Anatomy, School of Biomedical Sciences, University of Otago, P.O. Box 56, Dunedin, New Zealand.

## Abstract

RNA sequencing offers unprecedented access to the transcriptome. Key to this is the identification and quantification of many different species of RNA from the same sample at the same time. In this study we describe a novel protocol for simultaneous detection of coding and non-coding transcripts using modifications to the Ion Total RNA-Seq kit v2 protocol, with integration of QIASeq FastSelect rRNA removal kit. We report highly consistent sequencing libraries can be produced from both frozen high integrity mouse hippocampal tissue and the more challenging post-mortem human tissue. Removal of rRNA using FastSelect was highly efficient, resulting in less than 1.5% rRNA content in the final library, significantly better than other reported rRNA removal techniques. We identified >30,000 unique transcripts from all samples, including protein-coding genes and many unique species of non-coding RNA, in biologically-relevant proportions. Furthermore, normalized sequencing read count for select genes significantly negatively correlated with Ct values from RT-qPCR analysis from the same samples. These results indicate that this protocol accurately and consistently identifies and quantifies a wide variety of transcripts simultaneously. The highly efficient rRNA depletion, coupled with minimized sample handling and without complicated and high-loss size selection protocols, makes this protocol useful to researchers wishing to investigate whole transcriptomes.

## Introduction

Over the last 50 years, it has gradually been accepted that previously-dubbed “junk DNA” plays vital biochemical roles in higher organisms. This DNA does not directly code for proteins, yet makes up ~80% of the human genome. The gathering consensus is that by taking an holistic approach to the genome, not just examining protein-coding genes, it is possible gain a better understanding of the whole (1). This concept extends to the investigation of the transcriptome by RNA sequencing (RNA-Seq), which is already moving away from simply examining differential gene expression (DGE) of messenger RNA (mRNA), and towards including other species of cellular RNA. Many of these other classes of RNA are increasingly being shown to have numerous and varied biological roles, as well as being implicated in disease aetiology and pathogenesis.

Non-coding RNA (ncRNA) are attractive targets of research due to their stability (2–5), functional relevance to health and disease (6,7), and high level of evolutionary conservation (8), however, capturing these other RNA species can be more challenging than capturing mRNA. Many commercially available ncRNA sequencing kits exist, including but not limited to Illumina TruSeq Small RNA Sample kit, PerkinElmer NEXTFLEX Small RNA Library Prep Kit, and NEBNext Small RNA-Seq Kit. All of these methods rely on specifically isolating small RNA transcripts (usually <160bp) by size selection and excision of the region of interest from a solid matrix, followed by precipitating the RNA (9). While this can allow deep sequencing of RNA within that size range, it is limited in two respects. First, a significant amount of information on the transcriptome is lost through RNA falling outside of the excised size range. Second, the precipitation of RNA from the gel will never be entirely efficient, resulting in unavoidable loss of material. It would be ideal, therefore, to develop a protocol that allows the researcher to identify non-coding RNA transcripts, as well as larger coding- and non-coding RNA, without material loss due to size selection.

The removal of ribosomal RNA (rRNA) from RNA samples is a crucial step in RNA-Seq methods. Ribosomal RNA is a considerable roadblock to the detection of other functionally relevant RNA species, as it makes up to 80-90% of total RNA in a cell (by mass) (10–12). Current RNA-Seq protocols generally follow one of two rRNA removal methods – enrichment of polyadenylated (poly-A) RNA or depletion of ribosomal RNA (rRNA). Poly-A selection relies on the use of oligo dT primers to capture polyadenylated transcripts. This population is largely made up of mRNA, but does not capture all mRNA. Indeed, there is considerable evidence that a significant proportion of brain-derived mRNA is non-polyadenylated, further complicating the use of poly-A selection for investigating brain transcriptomes (13–17). As a result, RNA-Seq data generated by positive capture of polyadenylated RNA do not represent information from non-polyadenylated transcripts, degraded RNA transcripts, and the vast majority of non-coding RNA species. By contrast, depleting total RNA samples of rRNA allows quantification of a more varied population of RNA species. rRNA depletion can be achieved by a variety of means, including dedicated rRNA removal kits. For example, Ribo-Zero Plus (Illumina), captures rRNA by hybridization to complimentary oligonucleotides (ONTs) coupled to magnetic beads that, when precipitated, remove the rRNA from the rest of the RNA. Another method relies on hybridizing rRNA to complementary DNA oligonucleotides. This is followed by RNAseH digestion of the RNA:DNA hybrids (NEBNext rRNA Depletion kit, Takara Bio RiboGone). Takara/Clontech SMARTer Stranded Total RNA-Seq kit also includes a proprietary method for rRNA removal that uses ZapR to degrade cDNA originating from rRNA. These methods show different rRNA depletion efficiency, depending on input RNA quality (18–21) and furthermore, some variability in rRNA depletion efficiency has been reported between the implementation of the same protocol at different physical locations (18).

Generally, the literature reports rRNA making up anywhere from 0.5 − 20% of final rRNA-depleted sequencing libraries (18–22). With sequencing protocols usually generating in the vicinity of 20-30 million reads per sample, this can equate to 4-6 million reads mapping to rRNA. Better rRNA removal efficiency would result in those reads becoming available for mRNA and non-coding RNA sequence reads, of greater experimental interest to researchers. Furthermore, many of these techniques include multi-step protocols, often requiring precipitation steps (in the case of bead-based systems) and/or digestion or degradation steps. This often results in the loss of RNA material through purification, precipitation, or digestion. Thus the ideal rRNA removal technology would minimise workflow steps, sample handling, and reduce loss of material from precipitation or purification steps.

In a new development, Qiagen has released the QIAseq FastSelect rRNA removal kit, which utilizes complementary ONTs that bind to rRNA and prevent their reverse transcription to cDNA. The two main draws of this technology are its seamless integration into existing library preparation protocols (a single pipetting step and 14 minute protocol), and the fact that it does not require any additional purification, precipitation, or enrichment steps, thereby minimizing sample loss. This considerable reduction in sample handling is key to accurate and efficient detection of especially low-abundance transcripts.

Another perceived hurdle in effective RNA-Seq is quality of the input RNA. While standardized methods exist for assessing RNA quality and the level of RNA degradation (most commonly RNA Integrity Number; RIN), there is no well-defined consensus on what constitutes a sample that is too degraded for RNA-Seq. Any cut-off for sample exclusion used in the literature is, therefore, almost entirely arbitrary (23,24). This is not to say, however, that difficulties do not exist when performing RNA-Seq using samples of lower quality (25–27). Firstly, as noted above, degraded RNA proscribes the use of Poly-A selection for rRNA removal, as the process of degradation renders poly-A selection inefficient, and introduces a strong 3’ gene end bias to sequenced reads (28,29). Second, studies report that RNA samples of low quality (such as those obtained from post-mortem human tissue, in particular after a long post-mortem interval >24 hours) consistently show decreased proportions of mappable reads and a perceived reduction in sample complexity, with fewer highly-expressed genes and an abundance of low-expression genes (26). Here we describe an end-to-end RNA-Seq workflow and analysis pipeline that addresses some of the shortcomings of currently available protocols, in particular in rRNA depletion, minimisation of sample loss, and handling of varying input RNA quality. We demonstrate that this protocol is capable of identifying and quantifying both coding and non-coding RNA simultaneously from both high-quality and degraded RNA samples.

## Materials and Methods

### Animal Studies

All animal use was compliant with the New Zealand Animal Welfare Act 1991, and performed under guidelines and approval of the University of Otago Animal Ethics Committee (approval number DET09/15). In this study we utilised a double transgenic model of Alzheimer’s disease (APPswe/PS1dE9, B6C3 background, hereafter referred to as APP/PS1) originally sourced from The Jackson Laboratory (https://www.jax.org/strain/004462) and maintained as a colony at the University of Otago breeding facility. All mice were genotyped for the presence of human exon-9-deleted variant PSEN1, which co-segregates with the APPswe gene, as previously described (30). Male transgenic (tg) and wild-type (wt) littermates at 15 months old (*n* = 4 per group) were anaesthetised with sodium pentobarbitol and the brains removed into ice cold artificial cerebrospinal fluid solution (aCSF; in mM: 124 NaCl, 3.2 KCl, 1.25 NaH_2_PO_4_, 26 NaHCO_3_, 2.5 CaCl_2_, 1.3 MgCl_2_, 10 d-glucose). The left hippocampus was dissected and snap-frozen on dry ice. All samples were stored at −80°C until used. RNA extracted from mouse hippocampi is henceforth referred to by the identifier “Sample #”.

### Human Studies

Use of human tissue was compliant with the New Zealand Health and Disability Ethics Committee (14/STH/20/AM07) and the Human Research Ethics Committee of The University of Melbourne (1545740) and the Victorian Institute of Forensic Medicine (EC 18-2019). Post-mortem middle temporal gyrus (MTG) samples were received from the Victorian Brain Bank (VBB). Age-matched healthy control brains (n = 4; 2 male, 2 female; age 80.5 ± 8.8) were defined as free from Alzheimer’s Disease (AD) lesions with numbers of plaques and tangles below the cut-off values for a neuropathological diagnosis of AD (NIA Reagan criteria). No other neurological diseases were present. Alzheimer’s disease brains (n = 3; 3 female; age 76.5 ± 7.7) met the standard criteria for Alzheimer’s disease neuropathological diagnosis. There were no significant differences between the ages of the two groups (T-test, p=0.55). Patient gender was self-reported. All samples were stored at −80°C until used. RNA extracted from human MTG samples is henceforth referred to by the identifier “Patient #”. Patient demographics and case information are available in Supplementary Table 1.

### RNA extraction

Total RNA was extracted from previously-frozen tissue using the mirVana™ PARIS™ RNA isolation kit (Invitrogen; Cat #AM1556), according to the manufacturer’s instructions. The concentration and purity were determined by both spectrophotometry (A260, A260/280 respectively; NanoDrop 1000 Spectrophotometer; NanoDrop Technologies, Waltham, MA) and capillary electrophoresis (RNA Integrity Number [RIN], RNA 6000 Nano chip, Cat #5067-1511; Agilent Bioanalyzer 2100, Agilent Technologies).

### Library preparation

Except where explicitly stated, all samples, regardless of species or group of origin, were treated identically. Sequencing libraries were prepared for Ion Proton using the Ion Total RNA-Seq kit v2 (Life Technologies; Cat #4479789) largely following manufacturer’s instructions. Total RNA (500ng) was used as input to the Ion Total RNA-Seq kit v2 (356ng input was used instead of 500ng for Patient 1 due to low RNA yield), to which was added 1µL of 1:100 ERCC Spike-In Mix 1 (Invitrogen; Cat #4456740), commonly employed to control for cross-sample variation in library preparation. RNA fragmentation by RNAse III was performed at 37°C. The fragmentation time was optimised to 8 min for the mouse RNA, and 1 minute for the human RNA. This will vary depending on quality and integrity of the input RNA material. The resulting fragmented RNA was purified using the Magnetic Bead Cleanup Module (Life Technologies; Cat #4475486), and purified RNA eluted in 13µL nuclease-free water.

Ligation of Ion adapters (Ion RNA-Seq Primer Set v2; Cat #4479789) was performed using 3µL of the eluted purified fragmented RNA, added to 2µL Ion Adapter Mix v2 and 3µL Hybridization solution, and incubated in a thermal cycler at 65°C for 10 min followed by 30°C for 5 min. To this hybridization reaction was added 10µL 2X Ligation Buffer and 2µL Ligation Enzyme Mix, and incubated at 16°C for 16 hours in a thermal cycler. Following ligation, reverse transcription (RT) and rRNA removal was performed simultaneously as follows. RT master mix was prepared on ice (per sample; 1µL nuclease-free water, 4µL 10X RT buffer, 2µL 2.5mM dNTP Mix, 8µL ion RT Primer v2, 1µL QIAseq FastSelect rRNA removal agent). QIAseq FastSelect rRNA removal agent (Qiagen, Cat #334386) consists of ONTs complementary to ribosomal RNA sequences. These ONTs, when bound to rRNA sequences, prevent reverse transcription. The master mix was added to the ligation reaction, and incubated at 70°C for 10 minutes, followed by a step-wise cooldown (2 min at 65°C, 2 min at 60°C, 2 min at 55°C, 5 min at 37°C, 5 min at 25°C, hold at 4°C). This step is necessary for the oligonucleotides in the FastSelect rRNA removal agent to bind rRNA fragments and prevent reverse transcription. Finally, 4µL 10X Superscript Enzyme Mix was added to each reaction and the reactions incubated at 42°C for 30 min.

The resulting cDNA was purified using the Magnetic Bead Cleanup Kit, and eluted in 12µL nuclease-free water. In order to amplify the cDNA, 6µL of this elution was added to a master mix of 45µL Platinum PCR Supermix, 1µL Ion Xpress 3’ Barcode Primer, and 1µL Ion Xpress RNA Barcode BC## (Life Technologies; Cat #4475485). This mixture was amplified in a thermal cycler for 14 cycles (Hold 2 min 94°C; Cycle 2x [94°C 30s; 50°C 30s; 68°C 30s]; Cycle 14x [94°C 30s; 62°C 30s; 68°C 30s]; Hold 5 min 68°C). The amplified cDNA was purified again using the Magnetic Bead Cleanup Kit, and analysed by capillary electrophoresis (High Sensitivity DNA chip, Cat #5067-4626; Agilent Bioanalyzer 2100).

### Sequencing on Ion Proton Platform

The prepared sequencing libraries were diluted to equimolar concentrations of 100 pmol/L for pooling. Emulsion PCR was performed with the Ion OneTouch™ 2 system (Invitrogen) using the Ion PI™ Hi-Q™ OT2 200 kit (Invitrogen; Cat #A26434) according to the manufacturer’s instructions. The four pairs of mouse samples (tg and wt) were processed simultaneously end-to-end, as were the seven human MTG samples. Libraries were sequenced on Ion PI™ v3 chips (Invitrogen; Cat #A26770), prepared using the Ion PI™ HiQ™ Seq 200 kit (Invitrogen; Cat #A26433, A26772). The mouse samples of two pools of mixed barcoded libraries were sequenced on two Ion PI v3 chips (2 wt and 2 tg per chip), avoiding the use of all sequential barcodes on the same chip. Similarly, human MTG libraries were sequenced in two pools of mixed barcoded libraries – one contained a pool of four samples (2 AD and 2 control), the other a pool of three samples (1 AD and 2 control).

### Reverse Transcription Quantitative Polymerase Chain Reaction (qPCR)

For gene expression qPCR, using 350ng starting total RNA input from mouse samples, cDNA was generated using SuperScript IV First Strand Synthesis System (Invitrogen; Cat #18091050) per manufacturer’s instructions, utilizing priming by random hexamers. Of this cDNA, a 1:25 dilution was used for the qPCR reaction, which was performed using TaqMan Gene Expression Master Mix (Applied Biosystems, Cat #4369016), with the following TaqMan gene primers: Mouse *Hprt* (Assay ID: 03024075_m1), *Cst7* (00438351_m1), *Tyrobp* (00449152_m1), *c-Fos* (00487425_m1), *Trem2* (04209424_m1). The reactions were amplified on a Applied Biosystems ViiA 7 system as follows: Hold 50°C 2 minutes, hold 95°C 10 minutes, and 40 cycles at 95°C for 15 s and 60°C for 1 minute.

For miRNA, 10ng of total RNA from mouse samples was used. cDNA was generated using the TaqMan MicroRNA Reverse Transcription Kit (Applied Biosystems, Cat #4366596) according to manufacturer’s instructions. The qPCR reactions were prepared using TaqMan Universal PCR Master Mix (Applied Biosystems, Cat #4304437), with the following TaqMan miRNA primers: miR-34a-5p (Assay ID: 000426), miR-34c-5p (000428), miR-129-1-3p (002298), miR-210-3p (000512). The reactions were amplified on a Applied Biosystems VaaA 7 system as follows: Hold 50°C 2 minutes, hold 95°C 10 minutes, and 40 cycles at 95°C for 15 s and 60°C for 1 minute.

MicroRNA and mRNA qPCR data were processed separately to account for differing input RNA amounts, and in each case, raw Ct values were used for analysis.

### Data Analysis

Data from each barcoded library were separated into different data files automatically on the Ion Torrent Suite version 5.4 (life Technologies, USA). The Ion Torrent Suite was also used for analysis of ERCC Spike-In controls. Sequence read quality was evaluated using FastQC v0.11.5 (https://www.bioinformatics.babraham.ac.uk/projects/fastqc/) (31). Adapter sequences were trimmed from reads using AdapterRemoval v2.1.7 (https://github.com/MikkelSchubert/adapterremoval/) (32). Reads were then trimmed for quality using Trimmomatic v0.38 (http://www.usadellab.org/cms/?page=trimmomatic) (33) using a 5-base sliding window, cutting when the average quality per base drops below 20, and dropping reads less than 17 bases long.

Mouse RNA reads were aligned to the *Mus musculus* GRCm38.95 reference genome (available on the Ensembl website: http://www.ensembl.org/info/data/ftp/index.html) and human RNA reads to the *Homo sapiens* GRCh38.96 reference genome using STAR v2.5.4b (https://github.com/alexdobin/STAR) (34). Reference .gtf files for RNA biotypes (protein-coding, pseudogenes, snRNA, snoRNA, unknown [TEC], Mt-RNA, lncRNA, lincRNA, antisense) were extracted from the *Mus musculus* GRCm38.95 and *Homo sapiens* GRCh38.96 annotation files using the grep command.

MicroRNA (miRNA) were quantified from aligned counts using miRDeep2 v0.1.2 (https://github.com/rajewsky-lab/mirdeep2) (35).

Piwi-interacting RNA (piRNA) sequences were obtained from piRNABank (http://pirnabank.ibab.ac.in) (36).

Sequences for tRNA were obtained from the UCSC Genome Browser (http://genome.ucsc.edu).

Data were analysed using R version 4.0.2 in RStudio v1.3.959. The following packages were used: *edgeR* (37), *Rsubread* (38), *Rsamtools* (39), *stringr* (40), *ggplot2* (41), *matrixStats* (42), *pheatmap* (43), *tidyverse* (44). Additional statistics (regression/correlation) were also performed using R. Additional analysis and data visualisation performed using SeqMonk v1.45.1 (https://www.bioinformatics.babraham.ac.uk/projects/seqmonk/).

### Data Availability

The data discussed in this publication have been deposited in NCBI’s Gene Expression Omnibus (45) and are accessible through GEO Series accession numbers: GSE163877 (https://www.ncbi.nlm.nih.gov/geo/query/acc.cgi?acc=GSE163877) and GSE163878 (https://www.ncbi.nlm.nih.gov/geo/query/acc.cgi?acc=GSE163878).

## Results and Discussion

### Modified library preparation protocol consistently produces high quality sequencing libraries

To determine whether this protocol (overview shown in Figure 1) can effectively be used to create whole transcriptome sequencing libraries from total RNA, we extracted RNA from mouse hippocampal tissue using the mirVANA Paris kit (Invitrogen) total RNA procedure. The extracted RNA was consistently highly concentrated and of high quality, as reported by the Agilent Bioanalyzer RNA Nano Chip (Table 1). The RIN was 9.1 ± 0.05 (mean ± standard deviation), with an average concentration of 174.6 ± 27.7ng/µL. A260/280 ratios, as calculated by spectrometry, were > 2.12 (Table 1; 2.14 ± 0.01), strongly implying the lack of dsDNA contamination in the sample. Representative electropherograms of total input RNA, fragmented RNA, and amplified final libraries are shown in Figure 2. Total RNA (Fig. 2a) shows clear 18S and 28S rRNA peaks, while post-fragmentation these peaks become distributed with the overall RNA size distribution shifting downwards (Fig. 2b). The characteristic small RNA peak at ~100 nucleotides (nt) is also clearly seen, and is retained post-fragmentation. Figure 2c shows the size distribution of the final library.

**Table 1:** RNA extraction from mouse tissue resulted in high-integrity, highly-concentrated total RNA.

| Sample | RNA Integrity Number (RIN) | Concentration (ng/μL) | rRNA ratio [28s/18s] | Ratio A260/280 |
| --- | --- | --- | --- | --- |
| Sample 1 | 9 | 198 | 1.4 | 2.14 |
| Sample 2 | 9.1 | 188 | 1.5 | 2.13 |
| Sample 3 | 9.2 | 226 | 1.9 | 2.14 |
| Sample 4 | 9.4 | 167 | 1.9 | 2.16 |
| Sample 5 | 8.9 | 144 | 1.5 | 2.16 |
| Sample 6 | 9 | 147 | 1.5 | 2.15 |
| Sample 7 | 9.1 | 165 | 1.8 | 2.12 |
| Sample 8 | 9.1 | 162 | 1.8 | 2.15 |
| *Average ± SD* | $9.1 \pm 0.05$ | $174.6 \pm 27.7$ | $1.66 \pm 0.19$ | $2.14 \pm 0.01$ |

**Figure 1:**
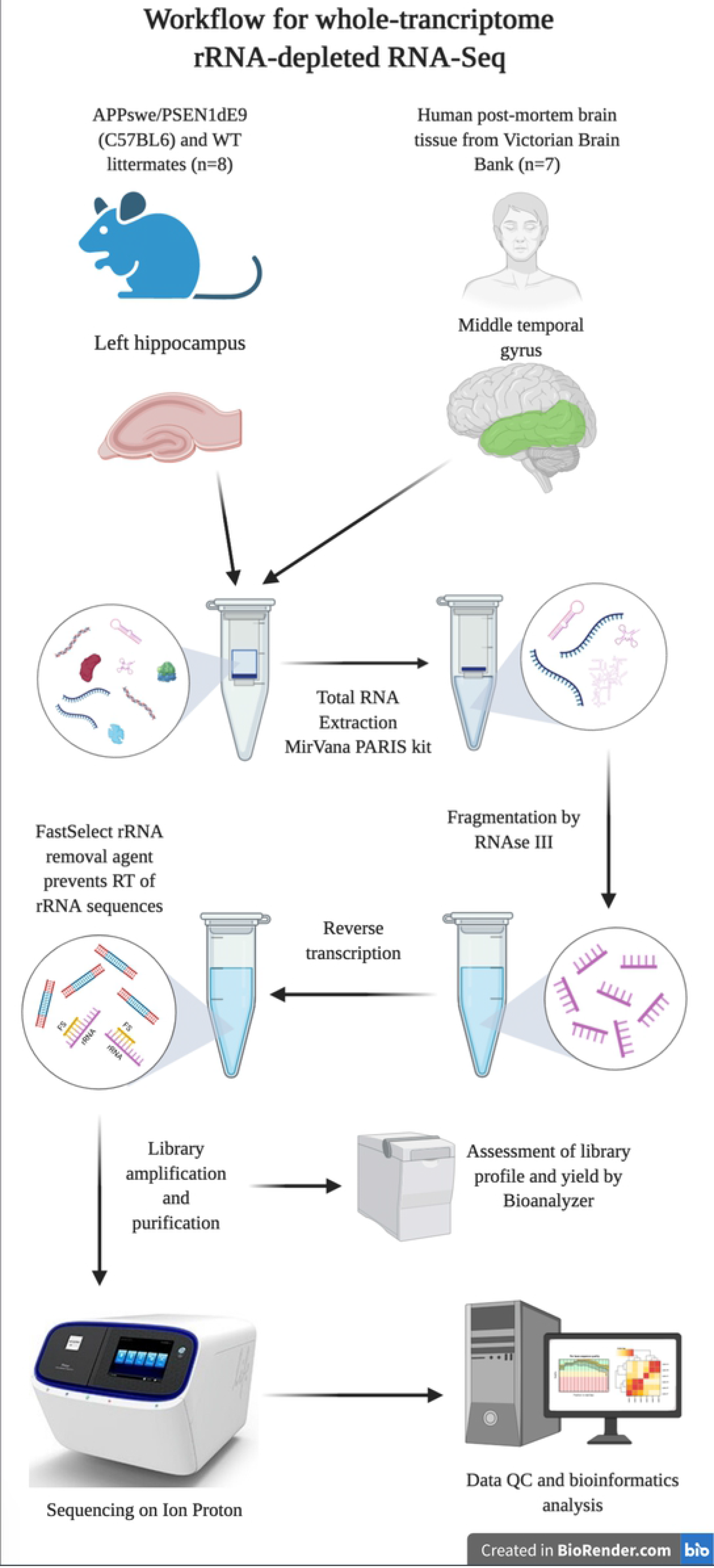
An overview of the protocol described here for simultaneous detection of coding and non-coding RNA by RNA-Seq. Created in BioRender.com.

**Figure 2:**
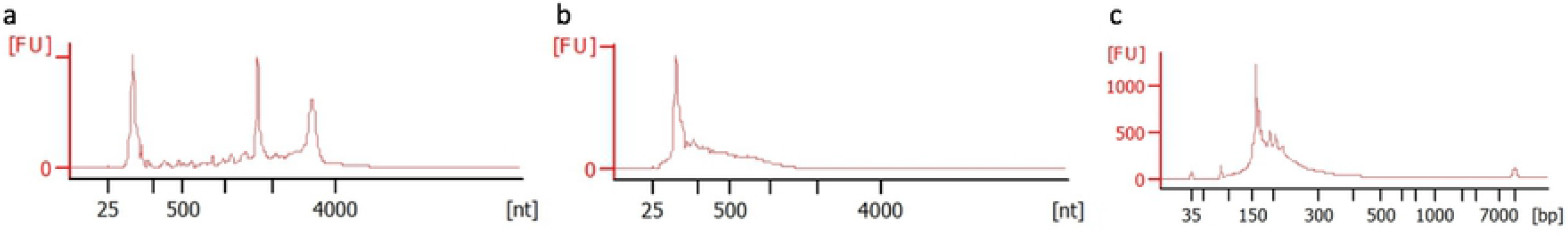
**Representative Bioanalyzer electropherograms of (a) total input RNA, (b) RNA after 8 minute fragmentation by RNAse III, and (c) final amplified cDNA library from mouse hippocampal RNA. Note the clear 18S and 28S rRNA peaks in total RNA (a) and their disappearance in fragmented RNA (b), as well as the downward shift of size distribution between total RNA (a), fragmented RNA (b) and final cDNA library (c).**

Next, we assessed the remainder of the library preparation protocol. We adjusted the adapter ligation period to a 16-hour (overnight) incubation at 16°C, rather than the recommended 30°C for 30 minutes. This markedly increased adapter ligation efficiency 15-fold (From 53 pg/µL to 745.5 pg/µL; Suppl. Figure 1). While this does, of course, increase the time required for library preparation, this significant increase in ligation efficiency outweighs this drawback.

Removal of rRNA was performed by addition of the Qiagen FastSelect rRNA removal agent to the cDNA synthesis steps. Hybridization of the FastSelect ONTs was achieved by the step-wise cooldown of the reaction mix from 65°C to 25°C, before addition of Superscript Enzyme Mix. The final cDNA libraries showed size distributions in-line with manufacturer’s recommendations (up to 200-base fragments for the Ion Proton system), with <50% of cDNA fragments falling under 160 base pairs (bp) (Fig. 2c). Since the libraries include small RNA species like miRNA which, including adapters, lie in the 40-45 bp size range, we expect to see a slightly higher proportion of the final library below 160 bp.

The library preparation protocol described here resulted in sequencing libraries ranging from 25.5 to 159.2 nmol/L in concentration. Library and sequence run metrics are given in Table 2.

**Table 2:** Library and sequencing metrics for mouse RNA samples.

| Sample | Final Library molarity (pmol/L) | Barcode ## | # Raw reads | Mean read length (bp) | # quality filtered reads | % Post-filtering |
| --- | --- | --- | --- | --- | --- | --- |
| Sample 1 | 25,963.90 | 1 | 17,487,705 | 63 | 11,991,333 | 73.36 |
| Sample 2 | 25,459.80 | 2 | 20,424,675 | 86 | 14,329,078 | 80.61 |
| Sample 3 | 71,161.70 | 5 | 31,493,446 | 92 | 22,422,158 | 77.9 |
| Sample 4 | 39,349.50 | 6 | 27,138,089 | 93 | 18,407,710 | 72.99 |
| Sample 5 | 109,683.40 | 3 | 21,131,492 | 96 | 16,486,584 | 83.77 |
| Sample 6 | 55,390.00 | 4 | 24,032,524 | 113 | 18,236,544 | 80.07 |
| Sample 7 | 90,893.50 | 7 | 25,463,277 | 90 | 19,414,166 | 80.23 |
| Sample 8 | 159,175.50 | 8 | 22,395,347 | 103 | 17,274,480 | 80 |

The mean read length for each library ranged from 63 to 113 bp, with an average of 23,695,819 total raw reads. Filtering of low quality reads using the specifications described in the methods and removal of adapter sequences resulted in between 11,991,333 (sample 1) and 22,422,158 reads (sample 3). Between 73.36% and 83.77% of raw reads remained post-processing. There was no significant correlation between either raw reads (R^2^ = 0.40) or filtered reads (R^2^ = 0.19) and input RNA RIN.

Alignment to the reference genome resulted in between 58% and 75% uniquely mapped reads, 17.5% and 35% multi-mapped reads, 1.33% and 2% mapped to too many (>10) loci, and 5-12% unmapped reads (Table 3/Figure 3). These figures are consistently on the higher end of previously reported mapping statistics (46).

**Table 3:**
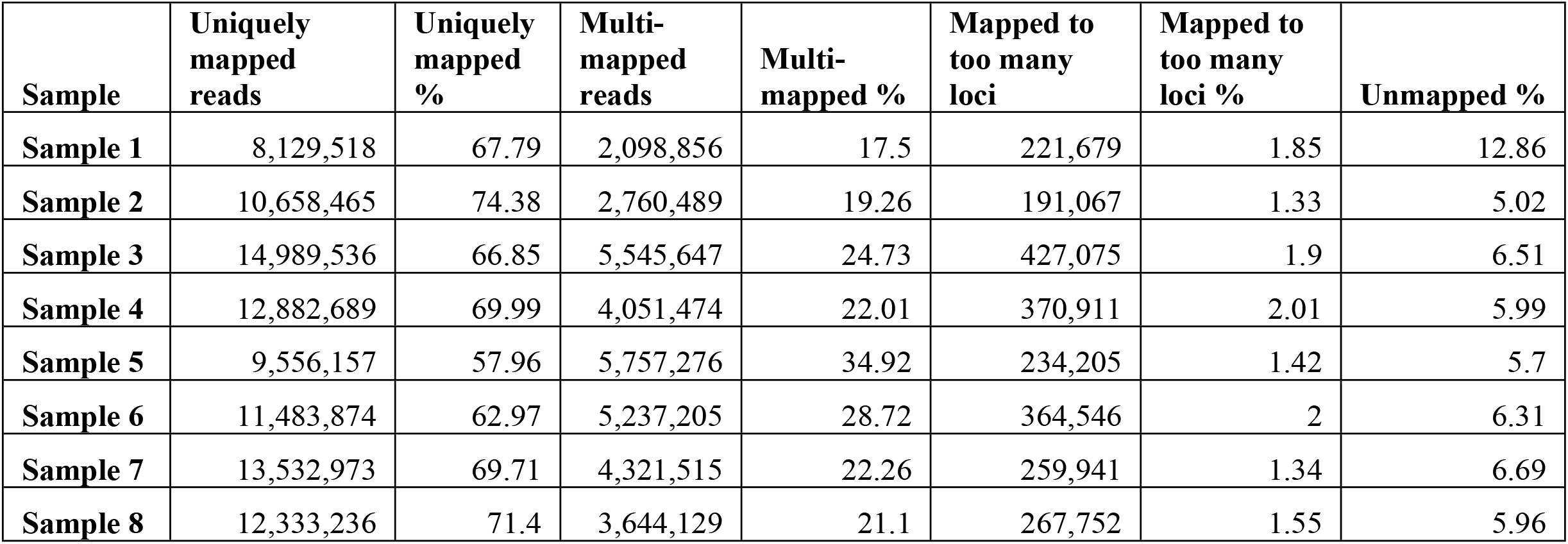
Read Alignment statistics for mouse RNA samples.

| Sample | Uniquely mapped reads | Uniquely mapped % | Multi-mapped reads | Multi-mapped % | Mapped to too many loci | Mapped to too many loci % | Unmapped % |
| --- | --- | --- | --- | --- | --- | --- | --- |
| Sample 1 | 8,129,518 | 67.79 | 2,098,856 | 17.5 | 221,679 | 1.85 | 12.86 |
| Sample 2 | 10,658,465 | 74.38 | 2,760,489 | 19.26 | 191,067 | 1.33 | 5.02 |
| Sample 3 | 14,989,536 | 66.85 | 5,545,647 | 24.73 | 427,075 | 1.9 | 6.51 |
| Sample 4 | 12,882,689 | 69.99 | 4,051,474 | 22.01 | 370,911 | 2.01 | 5.99 |
| Sample 5 | 9,556,157 | 57.96 | 5,757,276 | 34.92 | 234,205 | 1.42 | 5.7 |
| Sample 6 | 11,483,874 | 62.97 | 5,237,205 | 28.72 | 364,546 | 2 | 6.31 |
| Sample 7 | 13,532,973 | 69.71 | 4,321,515 | 22.26 | 259,941 | 1.34 | 6.69 |
| Sample 8 | 12,333,236 | 71.4 | 3,644,129 | 21.1 | 267,752 | 1.55 | 5.96 |

**Figure 3:**
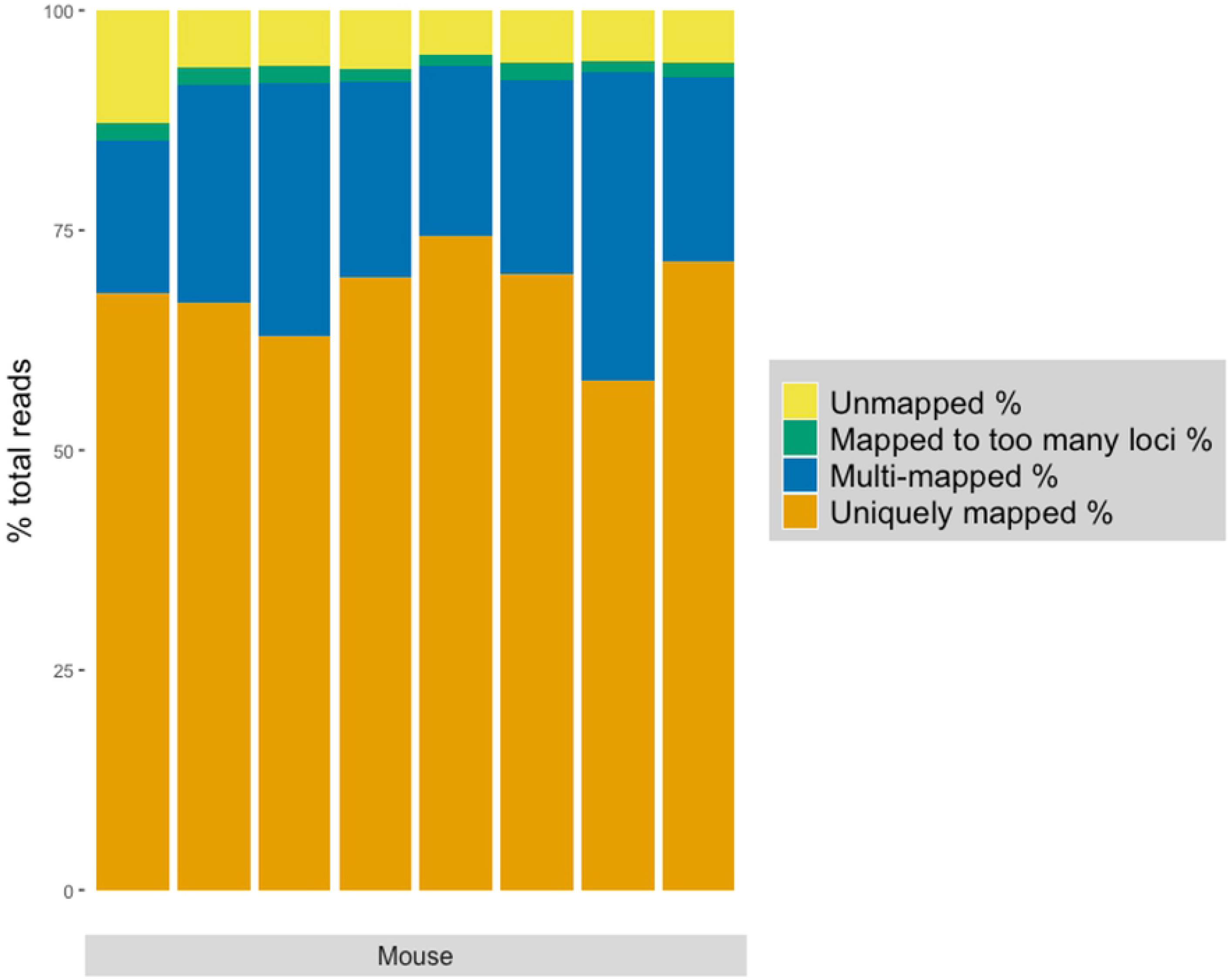
**RNA reads were mapped to the ENSEMBL *Mus musculus* GRCm38.95 annotated genome. Uniquely mapped reads, multi-mapped reads, reads mapped to too many loci (>10), and unmapped reads for each sample shown as a percentage of total trimmed and filtered reads.**

### High-quality sequencing libraries even from fresh-frozen human post-mortem brain tissue

RNA extracted from VBB brain tissue was less intact, as determined by electropherography, with an average RIN of 2.3 ± 0.2, and of lower concentrations than mouse RNA from similar amount of tissue input (68.57ng/µL ± 15.77; Table 4, Figure 4). A260/280 ratios, as calculated by NanoDrop, all lay above 2 (Table 4; 2.09 ± 0.04), again suggesting that the samples did not contain dsDNA. Notably, however, the resulting libraries were comparable in concentration and size distribution to those resulting from high quality mouse RNA (122.4 nmol/L ± 21.5; Table 5). Representative electropherograms of starting input RNA, fragmented RNA, and final libraries are shown in Figure 4. While the input RNA (Fig. 4a) lacks the defined 18S/28S rRNA peaks seen in Figure 2a, the 1 minute fragmented RNA (Fig. 4b) electropherogram shows a very similar size distribution to 8 minute fragmented mouse RNA (Fig. 2b). Again, the characteristic small RNA peak at ~100 nt is also clearly seen, and is retained post-fragmentation. Similarly, the final library size distribution (Fig. 4c) is comparable to that seen in Figure 2c. Library and sequencing statistics are shown in Table 5. The mean read length varied from 71 to 115 bp, with an average of 25,441,497 reads per sample. Quality filtering and adapter removal resulted in on average 15,855,518 reads per sample, leaving between 70% and 81% of reads post-processing. Again, there was no significant correlation between either raw reads (R^2^ = 0.005) or filtered reads (R^2^ = 0.0004) and input RNA RIN.

**Table 4.** RNA extraction from human post-mortem tissue resulted in low-integrity total RNA.

| Sample | RNA Integrity | Concentration | rRNA ratio | Ratio |
| --- | --- | --- | --- | --- |

| | Number (RIN) | (ng/μL) | (28s/18s) | A260/280 |
| --- | --- | --- | --- | --- |
| Patient 1 | 2.5 | 39.6 | 0.2 | 2.1 |
| Patient 2 | 1.9 | 84.2 | 0 | 2.11 |
| Patient 3 | 2.3 | 79.6 | 2.4 | 2.12 |
| Patient 4 | 2.2 | 80.9 | 0 | 2.02 |
| Patient 5 | 2.3 | 66.9 | 0.2 | 2.12 |
| Patient 6 | 2.5 | 57.4 | 0 | 2.05 |
| Patient 7 | 2.2 | 71.4 | 0 | 2.09 |
| Average ± SD | 2.27 ± 0.2 | 68.6 ± 15.8 | 0.4 ± 0.89 | 2.09 ± 0.04 |

**Table 5:** Library and sequencing statistics for human-derived RNA samples.

| Sample | Final Library molarity (pmol/L) | Barcode ## | # Raw reads | Mean read length | # quality filtered reads | % Post-filtering |
| --- | --- | --- | --- | --- | --- | --- |
| Patient 1 | 131,574.30 | 6 | 28,212,506 | 105 | 18,562,867 | 70.13 |
| Patient 2 | 131,068.10 | 2 | 28,789,188 | 109 | 19,003,266 | 73.98 |
| Patient 3 | 88,007.30 | 4 | 29,588,767 | 71 | 19,774,398 | 73 |
| Patient 4 | 134,256.40 | 3 | 20,982,426 | 106 | 15,431,060 | 81.11 |
| Patient 5 | 142,720.10 | 7 | 26,888,887 | 96 | 20,344,613 | 79.63 |
| Patient 6 | 134,278.50 | 1 | 23,085,633 | 101 | 16,661,101 | 79.13 |
| Patient 7 | 95,156.50 | 5 | 20,543,075 | 115 | 15,211,322 | 79.54 |

**Figure 4:**
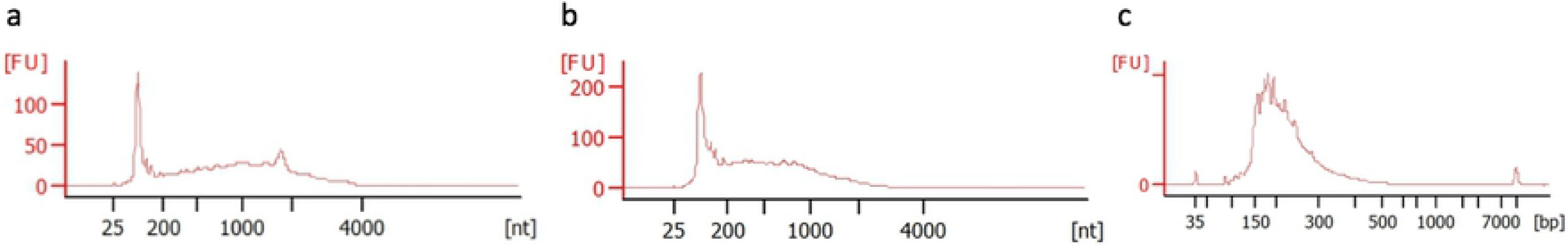
**Representative Bioanalyzer electropherograms of (a) total input RNA, (b) RNA after 1 minute fragmentation by RNAse III, and (c) final amplified cDNA library from human MTG RNA. Despite the lack of a distinct 18S/28S rRNA peak profile (a), fragmentation of RNA by RNAse III for 1 minute (b) resulted in a size distribution consistent with both previous preparations, and those recommended in the Ion Total RNA-Seq kit v2 User Guide. Final libraries (c) also show size distribution consistent with manufacturers recommendation, with <30% falling between 50 and 160 bp.**

Alignment to the human reference genome uniquely mapped between 79% and 82% of reads, and multi-mapped between 12% and 15% of reads (Table 6/Figure 5). Only ~3% of reads were mapped to too many loci, and between 2-3% of reads were unmapped. The proportion of uniquely-mapped reads is consistent with previously described mapping statistics, though with a considerably lower percentage of unmapped reads (46,47). We therefore demonstrate that the described protocol produces quality libraries from even fresh-frozen human post-mortem input RNA.

**Table 6:** Read Alignment statistics for human-derived RNA samples.

| Sample | Uniquely mapped reads | Uniquely mapped % | Multi-mapped reads | Multi-mapped % | Mapped to too many loci | Mapped to too many loci % | Unmapped % |
| --- | --- | --- | --- | --- | --- | --- | --- |
| Patient 1 | 14,689,409 | 79.13 | 2,869,622 | 15.46 | 554,660 | 2.99 | 2.42 |
| Patient 2 | 15,312,048 | 80.58 | 2,673,472 | 14.07 | 523,535 | 2.75 | 2.6 |
| Patient 3 | 14,815,243 | 74.92 | 3,796,504 | 19.2 | 628,450 | 3.18 | 2.7 |
| Patient 4 | 12,681,322 | 82.18 | 1,948,225 | 12.63 | 465,695 | 3.02 | 2.17 |
| Patient 5 | 16,171,475 | 79.49 | 3,154,060 | 15.5 | 627,631 | 3.08 | 1.92 |
| Patient 6 | 13,619,025 | 81.74 | 2,197,338 | 13.19 | 505,216 | 3.03 | 2.04 |
| Patient 7 | 12,427,195 | 81.7 | 2,033,866 | 13.37 | 398,154 | 2.62 | 2.32 |

**Table 6:** **Percentage mouse RNA reads mapped to gene biotypes per sample, as annotated in the *Mus musculus* GRCm38.95, as well as tRNA annotations from UCSC Genome Browser, piRNA annotations from piRNABank.**

| Samples | lincRNA | snoRNA | snRNA | Pseudogenes | piRNA | miRNA | miscRNA | Antisense | Unknown | Protein coding | lncRNA | tRNA | rRNA | Mt-rRNA | Mt-tRNA |
| --- | --- | --- | --- | --- | --- | --- | --- | --- | --- | --- | --- | --- | --- | --- | --- |
| 1 | 2.65 | 15.03 | 6.74 | 3.15 | 1.94 | 3.71 | 4.38 | 1.20 | 1.33 | 51.62 | 0.03 | 3.80 | 2.52 | 0.53 | 1.38 |
| 2 | 2.13 | 7.13 | 11.09 | 3.27 | 3.02 | 2.35 | 10.07 | 1.06 | 0.48 | 54.91 | 0.04 | 1.72 | 2.14 | 0.20 | 0.40 |
| 3 | 1.31 | 2.72 | 18.03 | 4.12 | 2.63 | 4.65 | 13.77 | 0.79 | 0.39 | 47.92 | 0.03 | 0.87 | 1.70 | 0.25 | 0.81 |
| 4 | 1.51 | 2.54 | 4.11 | 4.50 | 1.72 | 7.62 | 20.06 | 0.96 | 0.52 | 52.07 | 0.04 | 0.59 | 2.83 | 0.31 | 0.61 |
| 5 | 1.32 | 3.73 | 18.61 | 2.90 | 2.92 | 4.14 | 17.15 | 0.77 | 0.42 | 44.04 | 0.03 | 0.99 | 1.03 | 0.16 | 1.79 |
| 6 | 1.15 | 2.10 | 23.01 | 2.45 | 4.05 | 5.95 | 21.93 | 0.66 | 0.36 | 36.58 | 0.03 | 0.81 | 0.23 | 0.13 | 0.57 |
| 7 | 1.71 | 3.25 | 17.23 | 3.60 | 3.63 | 3.41 | 9.95 | 1.02 | 0.60 | 50.99 | 0.04 | 0.66 | 2.58 | 0.38 | 0.98 |
| 8 | 1.55 | 3.10 | 16.55 | 3.53 | 3.57 | 3.34 | 10.76 | 0.94 | 0.45 | 53.68 | 0.04 | 0.72 | 1.15 | 0.17 | 0.46 |

**Figure 5:**
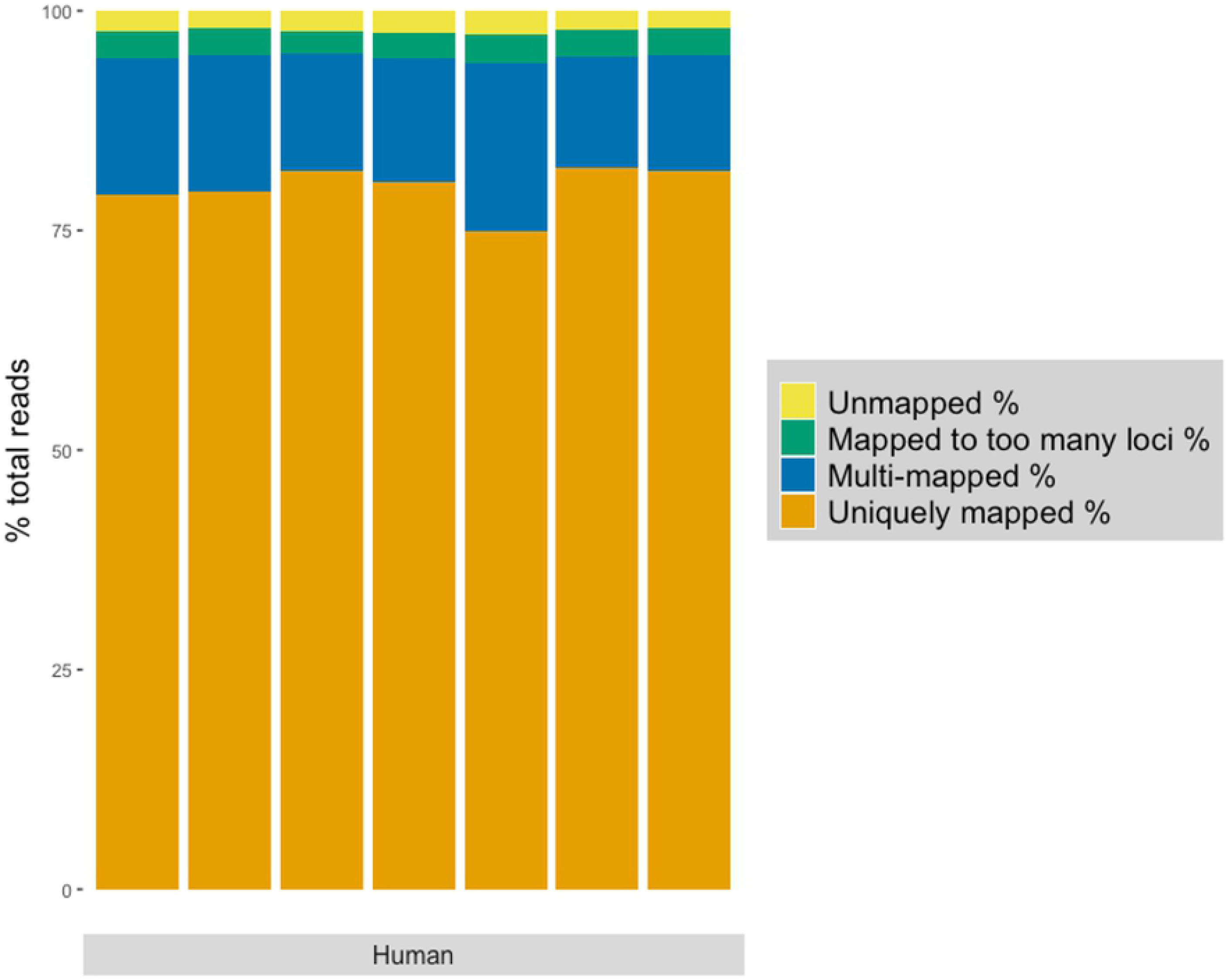
**RNA reads were mapped to the ENSEMBL *Homo sapiens* GRCh38.96 annotated genome. Uniquely mapped reads, multi-mapped reads, reads mapped to too many loci (>10), and unmapped reads for each sample shown as a percentage of total reads.**

### Qiagen FastSelect rRNA removal agent results in minimal rRNA content

To assess the effectiveness of Qiagen FastSelect rRNA removal agent, we used SeqMonk RNA-Seq QC to quantify the percentage of reads mapped to rRNA sequences in both the mouse and human samples. Ribosomal RNA content in RNA extracted from the mouse hippocampal tissue was between 0.23 – 1.24% (0.81 ± 0.37; n=8), and alignment to mitochondrial RNA (Mt-rRNA and Mt-tRNA) accounted for, on average, 0.25 ± 0.17% and 0.87 ± 0.49% respectively. Similarly, in the human RNA, the same protocol quantified rRNA content between 0.034 – 0.39% (0.11 ± 0.13; n=7), with mitochondrial rRNA and tRNA accounting for, on average, 0.28 ± 0.27 and 5.11 ± 3.23% respectively. As an average Ion PI Chip loaded with four samples returns ~25 million reads per sample, a total RNA library prep without rRNA depletion would result in ~22 million reads mapping to rRNA, whereas the protocol described here resulted in only ~100-200,000 reads mapped to rRNA. Compared to other techniques for rRNA removal from sequencing libraries, the technique described here performed consistently better that previously published methods, which range anywhere from 1% to 20% rRNA content (18–22). Thus the considerable reduction in rRNA content achieved by our protocol frees up valuable sequencing resources.

### This library preparation protocol and analysis pipeline identifies a variety of coding- and non-coding-RNA in biologically-relevant proportions

To achieve an estimate of the ability of this workflow to identify transcripts of interest, we performed bioinformatic analysis to determine a) how many different transcripts can be identified from the RNA-Seq data and b) what kind of transcripts can be identified. We calculated Reads Per Kilobase Million (RPKM) for mouse and human RNA samples to normalise the number of unique transcripts detected for sequencing depth and gene length. Mouse samples identified >18,500 unique transcripts expressed at greater than one RPKM and in total ~31,000 expressed at greater than 0.1 RPKM, representing ~50% of total annotated transcripts in the reference genome (Figure 6). For the human samples, a similar number of transcripts were found at >1 RPKM, with more than ~35,000 unique transcripts at >0.1 RPKM, representing ~60% of the total annotated transcripts in the reference genome (Figure 6). This is in stark contrast to some of the reported difficulties in RNA-Seq using low-quality input RNA – notably decreased proportions of mappable reads and reduction in sample complexity (26). In fact, our data suggest that this protocol results in proportions of successfully mapped reads and levels of gene expression comparable to high-quality, undegraded RNA samples.

**Figure 6:**
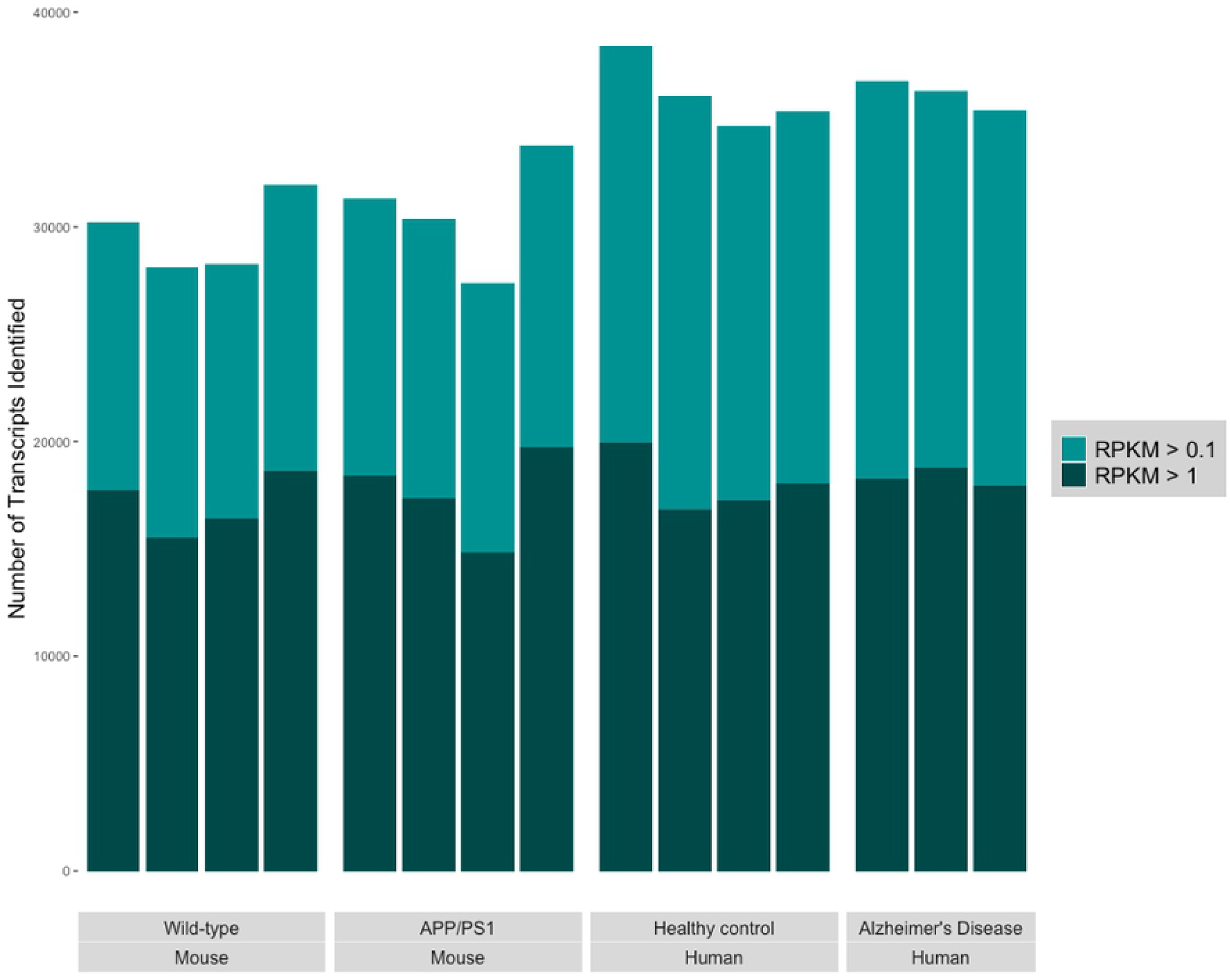
**Number of genes detected at >1 Reads per Kilobase Million (RPKM; dark green) and >0.1 RPKM (light green) for each mouse sample and human sample. RPKM here was used as a proxy for normalized expression. The number of genes detected at RPKM >0.1 is suggestive of the ability of this workflow to detect large numbers of low-expressed genes, many of which are non-coding.**

Breakdown of read alignment by transcript biotype (as annotated in each reference genome – *Mus musculus* GRCm38.100 and *Homo sapiens* GRCh38.96 – as well as piRNA and tRNA from piRNABank and UCSC Genome Browser respectively) is shown in Table 6 and 7. The average percentage content by gene biotype is shown in Figure 7. The largest number of reads mapped to protein-coding mRNA, (Mouse − 48.86% ± 6.02; human – 40.22% ± 4.43). There were numerous alignments to various species of ncRNA, including miRNA, piRNA, snRNA, snoRNA, lincRNA, and pseudogenes. As expected, a higher proportion of reads mapped to non-coding RNA in the human samples compared to the mouse, consistent with recent indications that organism complexity is reliant on non-coding RNA, rather than genome size (48). With the removal of rRNA from the prepared libraries, proportions of ncRNA correspond approximately to reported cellular RNA contents (49). Many species of non-coding RNA show very narrow variance between samples (Fig. 8), while others varied significantly. In particular, small nuclear RNA (snRNA) expression was highly variable, ranging from 4 to 23% in mouse RNA (Fig. 8a), and 2.5 to 15.5% in human RNA (Fig. 8b). Despite their frequent use as reference genes for qPCR and gene array experiments, individual variability in snRNA expression has been reported previously (50,51), and these observations are supported by the data presented here. One important caveat to consider, however, is that PCR amplification is known to favour smaller fragments over larger ones (52). As such, it is possible (even likely) that transcripts associated with small non-coding RNA are overrepresented in our final sequencing libraries. However, this effect is likely consistent across samples and libraries, and would certainly not affect the identification of unique transcripts. As is often recommended for RNA-Seq experiments, further investigation and validation of differentially-expressed transcripts by, for example, quantitative PCR would address this concern with regards to absolute quantification.

**Table 7:** **Percentage human RNA reads mapped to gene biotypes per sample, as annotated in the *Homo sapiens* GRCh38.96, as well as tRNA annotations from UCSC Genome Browser, piRNA annotations from piRNABank.**

| Samples | lincRNA | snoRNA | snRNA | Pseudogenes | piRNA | miRNA | Antisense | Unknown | Protein coding | lncRNA | tRNA | rRNA | Mt-rRNA | Mt-tRNA |
| --- | --- | --- | --- | --- | --- | --- | --- | --- | --- | --- | --- | --- | --- | --- |
| Patient 1 | 4.04 | 29.44 | 6.70 | 2.96 | 1.45 | 0.91 | 3.19 | 0.22 | 37.19 | 0.03 | 8.08 | 0.26 | 0.19 | 5.36 |
| Patient 2 | 4.23 | 19.54 | 6.53 | 4.26 | 2.49 | 0.75 | 3.74 | 0.27 | 46.67 | 0.03 | 9.33 | 0.46 | 0.23 | 1.46 |
| Patient 3 | 4.18 | 23.93 | 2.54 | 2.35 | 1.43 | 2.57 | 2.79 | 0.36 | 33.23 | 0.03 | 12.96 | 1.34 | 0.89 | 11.39 |
| Patient 4 | 5.15 | 22.74 | 10.61 | 2.73 | 1.58 | 1.06 | 2.36 | 0.39 | 42.79 | 0.03 | 4.10 | 0.32 | 0.20 | 5.96 |
| Patient 5 | 4.05 | 26.91 | 7.98 | 2.08 | 1.49 | 1.21 | 3.07 | 0.14 | 38.50 | 0.02 | 8.55 | 0.34 | 0.19 | 5.45 |
| Patient 6 | 4.43 | 22.71 | 15.49 | 2.49 | 1.35 | 0.73 | 2.78 | 0.28 | 40.04 | 0.03 | 5.72 | 0.24 | 0.13 | 3.60 |
| Patient 7 | 5.59 | 23.90 | 11.28 | 2.88 | 1.60 | 0.58 | 2.75 | 0.25 | 43.15 | 0.03 | 5.08 | 0.25 | 0.13 | 2.52 |

**Figure 7:**
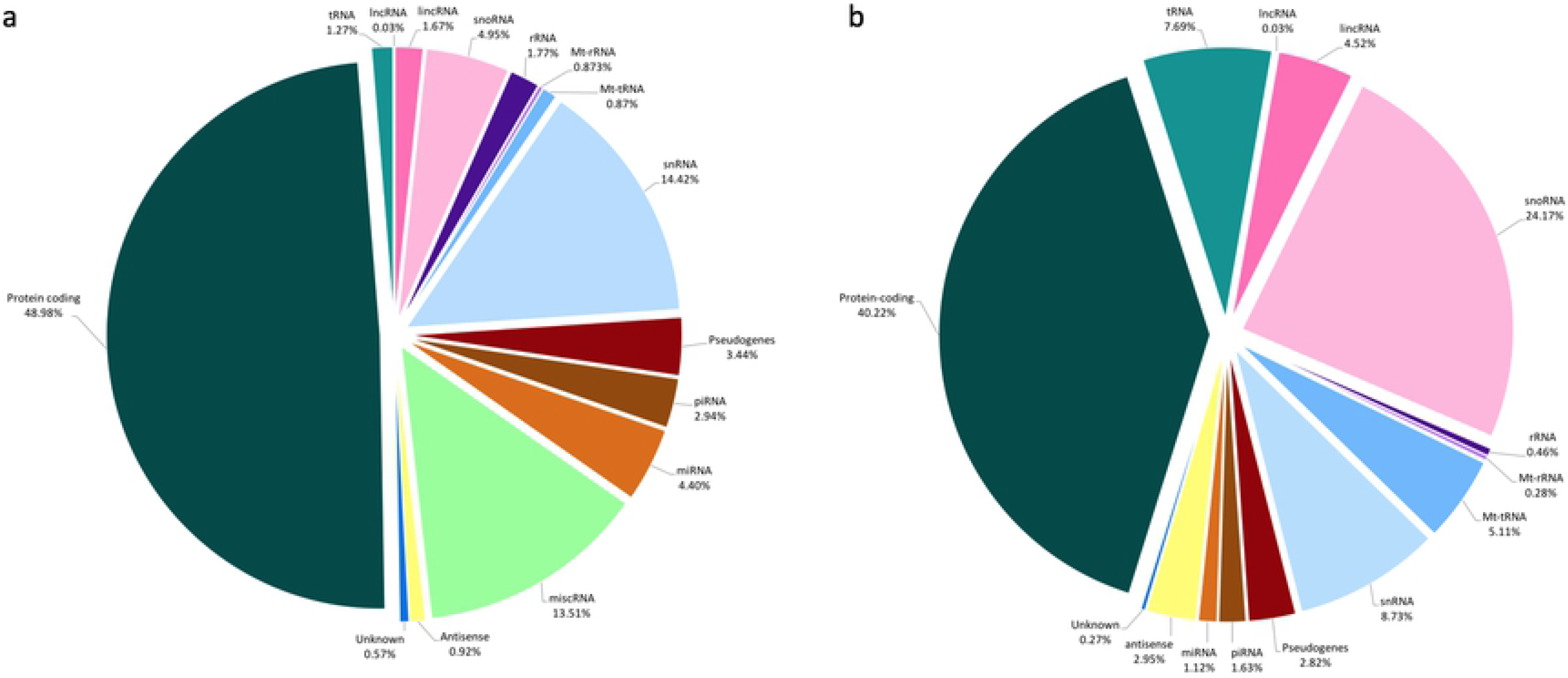
**Percentage RNA reads mapped to gene biotypes for (a) mouse and (b) human samples, averaged across samples. The largest proportion for both samples is made up of protein coding RNA. However in both mouse and human RNA, protein-coding RNA made up less than 50% of the total, with the majority being various forms of non-coding RNA.**

**Figure 8:**
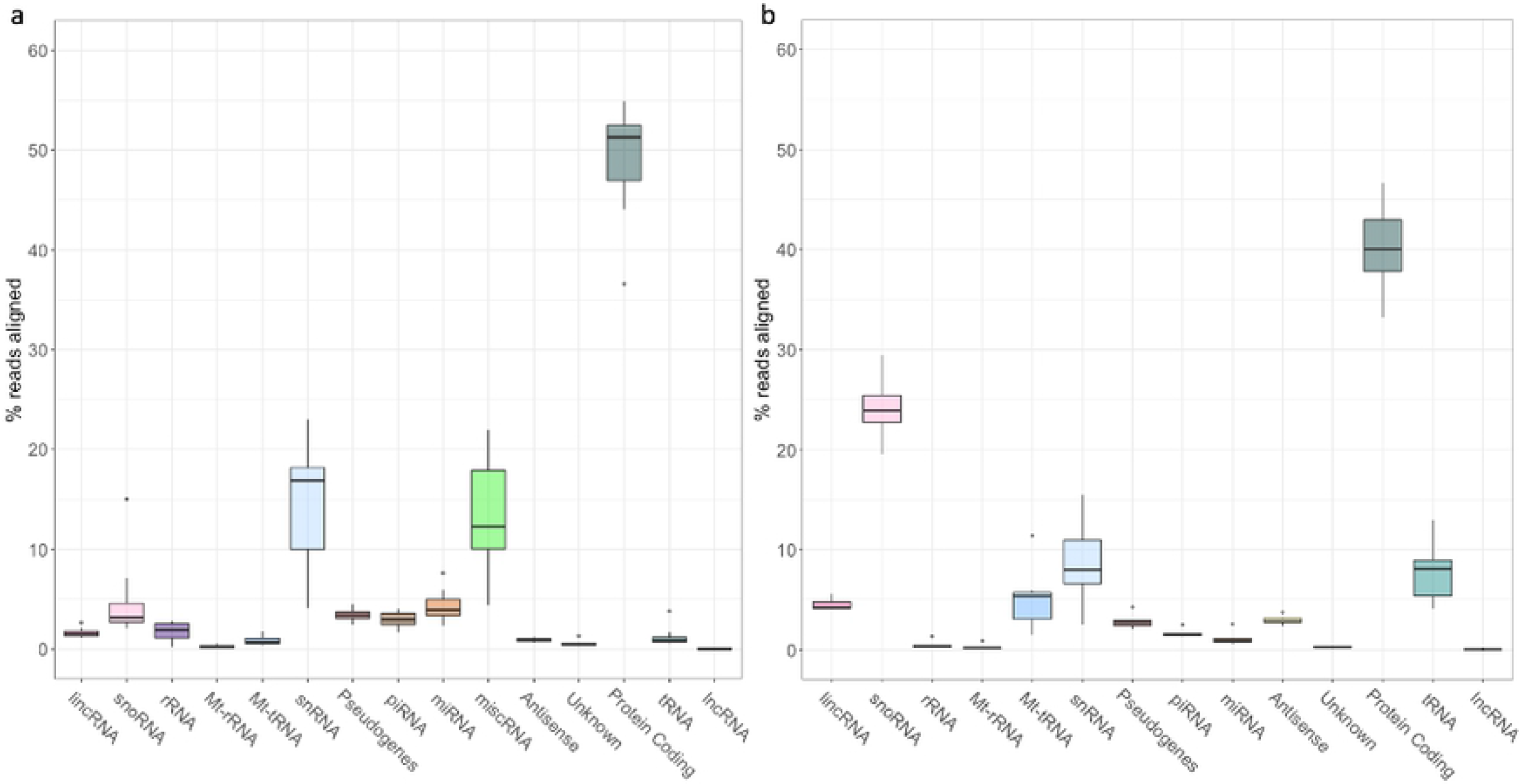
**Box and whisker plot showing the range of percentages of reads mapped to gene biotypes for (a) mouse and (b) human samples. The majority of RNA species in both samples show very small ranges, while some (notably snRNA and snoRNA) are more variable between samples.**

### Select gene sequencing reads significantly correlate with cycle threshold (Ct) values obtained by quantitative PCR

In order to ascertain to what extent sequencing reads obtained from this protocol can be representative of the actual number of RNA molecules in the sample, we performed RT-qPCR analysis of selected genes and miRNA from the mouse samples. We then determined the correlation coefficient (Pearson’s Product-Moment Correlation) of Ct values against RPKM, in order to control for library size and gene length (Figure 9). We found that both mRNA (*Hprt, Trem2, Tyrobp, c-Fos*, and *Cst7*; R^2^ = −0.81, p = 3.2e-10; Fig. 9a) and miRNA (miR-129-1-3p, miR-34a-5p, miR-34c-5p, miR-210-3p; R^2^ = −0.59, p = 0.00036; Fig. 9b) showed significant negative correlation between Ct values and normalized sequencing reads. This strongly suggests that data obtained from this sequencing method results in read counts that are largely representative of actual RNA content, and lends credence to differential gene expression analyses performed with these data.

**Figure 9:**
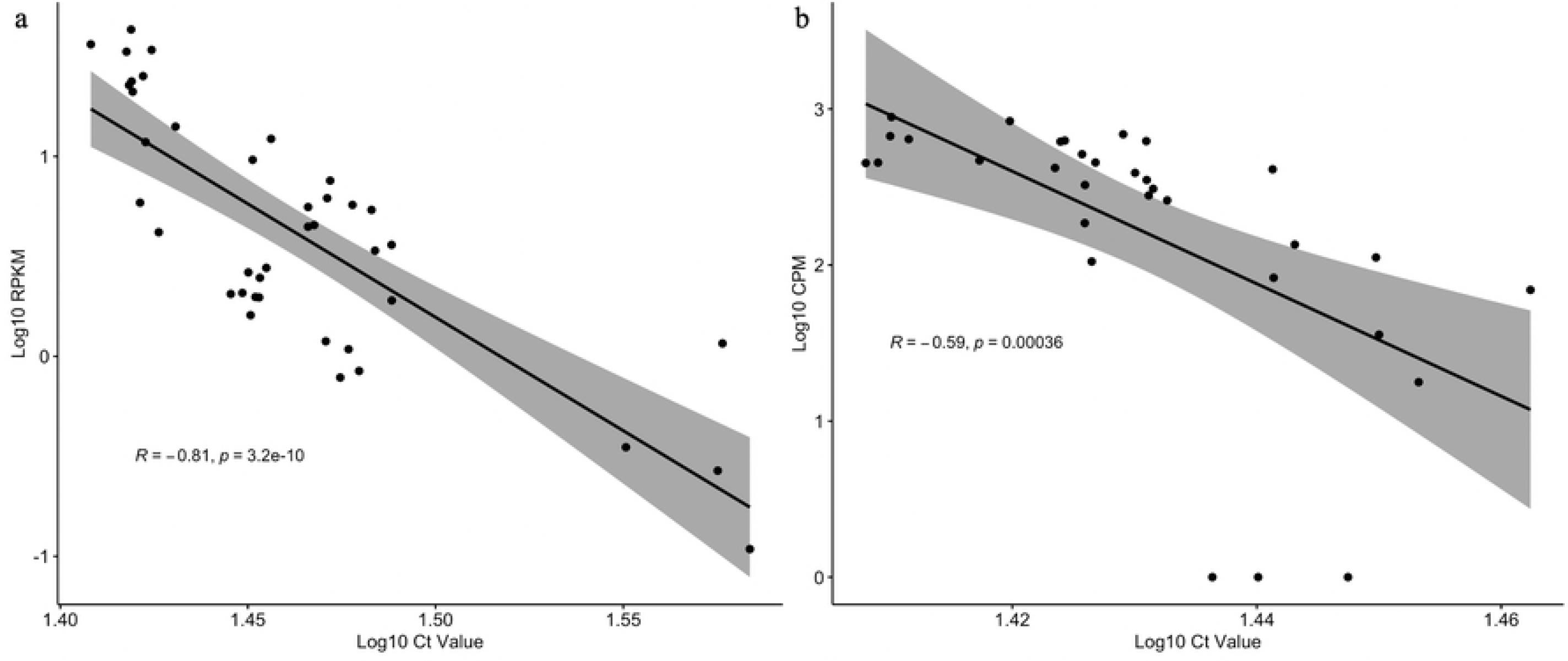
**Correlation between Ct values from qPCR and normalized sequencing reads mapped to select genes (a) and miRNA (b). Gene sequencing reads were normalized for library size and gene length using RPKM, while miRNA reads were normalized for library size using CPM. Both sets of data passed Shapiro-Wilk tests of normality (p>0.05). The linear regression line, confidence interval, Pearson’s correlation coefficient, and significance value are indicated.**

## Conclusions

RNA sequencing is an ever-evolving technique that offers unique insights into the transcriptome. Current protocols often require the researcher to choose between investigating mRNA (by poly-A selection) or small RNA (by size selection). Either one of these alone, while offering great depth of sequencing, misses out on a great deal of information from excluded transcripts. There is room, therefore, for a protocol to investigate the whole transcriptome, from the same sample at the same time.

Here we report a novel method for whole transcriptome ribosomal-depleted RNA-Seq. This approach takes advantage of several existing commercially available kits, with some important alterations to manufacturer’s protocols. This altered workflow resulted in high quality sequencing libraries from input RNA samples of a variety of quality, from both mouse and human tissue. Low quality input RNA had no negative effect on the final library quality. Qiagen FastSelect rRNA removal agent integrated seamlessly into the existing Ion Total RNA-Seq kit v2 library prep protocol, and resulted in highly effective depletion of rRNA from the final libraries, even from degraded samples, which is often a drawback of other rRNA removal techniques. A high number of genes were identified in the RNA-Seq data, including transcripts often overlooked by more targeted RNA-Seq protocols (refer Fig. 7). The majority of reads mapped to species of non-coding RNA, and most of these were also highly consistent between samples within each species. Furthermore, sequencing reads (normalized to library size and gene length) correlate significantly with Ct values from qPCR quantitation (which allows for the most precise quantification of RNA content), suggesting that read counts obtained from this RNA-Seq protocol can be used to infer quantitative gene expression.

A similar protocol (fragmented, ribodepleted TGIRT-Seq) has been previously reported (53–55), that also aimed to simultaneously sequence coding and non-coding RNA. While it has seen some limited implementation since (56,57), it is yet to be widely accepted. There are a few factors where we believe improvements can be made. First, the use of QIASeq FastSelect for rRNA depletion we found superior to the RiboZero Gold used by Boivin and colleagues (reagent now defunct). While we did not compare these two methods directly, the comparison can be inferred through the literature. Crucially, rRNA-removal by FastSelect requires significantly less sample handling than RiboZero Gold, and does not require additional bead purification, preventing sample loss. Second, the protocol described here appears to give more representative non-coding RNA reads – in particular with regards to miRNA and piRNA – as compared to estimated abundances reported in literature (49,53). Altogether, we believe this workflow may be useful to researchers wishing to investigate the whole transcriptome simultaneously, with effective rRNA depletion, and without complicated and high-loss size selection protocols commonly used for small RNA-Seq, or poly-A selection for mRNA-Seq.

## Acknowledgements

Human post-mortem tissues were received from the Victorian Brain Bank Network, supported by The Florey, The Alfred and the Victorian Institute of Forensic Medicine and funded in part by Parkinson’s Victoria, MND Victoria, Fight MND and Yulgilbar Foundation. We acknowledge Dr. Bruce Mockett for management of the transgenic mouse colony.

## References

1. Pennisi E. ENCODE Project Writes Eulogy for Junk DNA. Science (80-). 2012;337(6099):1159–61.

2. Ayupe AC, Tahira AC, Camargo L, Beckedorff FC, Verjovski-Almeida S, Reis EM. Global analysis of biogenesis, stability and sub-cellular localization of lncRNAs mapping to intragenic regions of the human genome. RNA Biol. 2015;12(8):877–92.

3. Clark MB, Johnston RL, Inostroza-Ponta M, Fox AH, Fortini E, Moscato P, et al. Genome-wide analysis of long noncoding RNA stability. Genome Res. 2012;22:885– 98.

4. Tani H, Mizutani R, Salam KA, Tano K, Ijiri K, Wakamatsu A, et al. Genome-wide determination of RNA stability reveals hundreds of short-lived noncoding transcripts in mammals. Genome Res. 2012;22:947–56.

5. Friedel CC, Dölken L, Ruzsics Z, Koszinowski UH, Zimmer R. Conserved principles of mammalian transcriptional regulation revealed by RNA half-life. Nucleic Acids Res. 2009;37(17):115.

6. Laura Idda M, Munk R, Abdelmohsen K, Gorospe M. Noncoding RNAs in Alzheimer’s Disease HHS Public Access. Wiley Interdiscip Rev RNA. 2018;9(2).

7. Schwarzenbach H, Nishida N, Calin GA, Pantel K. Clinical relevance of circulating cell-free microRNAs in cancer. Nat Rev | Clin Oncol. 2014;11:145–56.

8. Qu Z, Adelson DL. Evolutionary conservation and functional roles of ncRNA. Front Genet. 2012 Oct;3(OCT):205.

9. Yeri A, Courtright A, Danielson K, Hutchins E, Alsop E, Carlson E, et al. Evaluation of commercially available small RNASeq library preparation kits using low input RNA. BMC Genomics. 2018 May;19(1).

10. Duncan R, Hershey JW. Identification and quantitation of levels of protein synthesis initiation factors in crude HeLa cell lysates by two-dimensional polyacrylamide gelelectrophoresis. J Biol Chem. 1983 Jun;258(11):7228–35.

11. Wolf SF, Schlessinger D. Nuclear metabolism of ribosomal RNA in growing, methionine-limited, and ethionine-treated HeLa cells. Biochemistry. 1977 Jun;16(12):2783–91.

12. Blobel G, Potter VR. Studies on free and membrane-bound ribosomes in rat liver. I. Distribution as related to total cellular RNA. J Mol Biol. 1967 Jun;26(2):279–92.

13. Harris DA, Sherbany AA. Cloning of non-polyadenylated RNAs from rat brain. Mol Brain Res. 1991;10(1):83–90.

14. Van Ness J, Maxwell IH, Hahn WE. Complex population of nonpolyadenylated messenger RNA in mouse brain. Cell. 1979 Dec;18(4):1341–9.

15. Snider BJ, Morrison-Bogorad M. Brain non-adenylated mRNAs. Brain Res Rev. 1992;17(3):263–82.

16. McKernan KJ, Peckham HE, Costa GL, McLaughlin SF, Fu Y, Tsung EF, et al. Sequence and structural variation in a human genome uncovered by short-read, massively parallel ligation sequencing using two-base encoding. Genome Res. 2009 Sep;19(9):1527–41.

17. Yang L, Duff MO, Graveley BR, Carmichael GG, Chen L-L. Genomewide characterization of non-polyadenylated RNAs. Genome Biol. 2011;12(2):R16.

18. Herbert ZT, Kershner JP, Butty VL, Thimmapuram J, Choudhari S, Alekseyev YO, et al. Cross-site comparison of ribosomal depletion kits for Illumina RNAseq library construction. BMC Genomics. 2018;19:1–10.

19. Huang R, Jaritz M, Guenzl P, Vlatkovic I, Sommer A, Tamir IM, et al. An RNA-Seq Strategy to Detect the Complete Coding and Non-Coding Transcriptome Including Full-Length Imprinted Macro ncRNAs. PLoS One. 2011 Nov;6(11):e27288.

20. Culviner PH, Guegler CK, Laub MT. A Simple, Cost-Effective, and Robust Method for rRNA Depletion in RNA-Sequencing Studies. Cooper VS, editor. MBio. 2020 Apr;11(2):e00010–20.

21. Cui P, Lin Q, Ding F, Xin C, Gong W, Zhang L, et al. A comparison between ribo-minus RNA-sequencing and polyA-selected RNA-sequencing. Genomics. 2010;96(5):259–65.

22. Haile S, Corbett RD, Bilobram S, Mungall K, Grande BM, Kirk H, et al. Evaluation of protocols for rRNA depletion-based RNA sequencing of nanogram inputs of mammalian total RNA. PLoS One. 2019 Oct;14(10):e0224578.

23. Imbeaud S, Graudens E, Boulanger V, Barlet X, Zaborski P, Eveno E, et al. Towards standardization of RNA quality assessment using user-independent classifiers of microcapillary electrophoresis traces. Nucleic Acids Res. 2005 Mar;33(6):e56.

24. Weis S, Llenos IC, Dulay JR, Elashoff M, Martínez-Murillo F, Miller CL. Quality control for microarray analysis of human brain samples: The impact of postmortem factors, RNA characteristics, and histopathology. J Neurosci Methods. 2007 Sep;165(2):198–209.

25. Schuierer S, Carbone W, Knehr J, Petitjean V, Fernandez A, Sultan M, et al. A comprehensive assessment of RNA-seq protocols for degraded and low-quantity samples. BMC Genomics. 2017;18(1):442.

26. Gallego Romero I, Pai AA, Tung J, Gilad Y. RNA-seq: impact of RNA degradation on transcript quantification. BMC Biol. 2014;12(1):42.

27. Li S, Tighe SW, Nicolet CM, Grove D, Levy S, Farmerie W, et al. Multi-platform assessment of transcriptome profiling using RNA-seq in the ABRF next-generation sequencing study. Nat Biotechnol. 2014 Sep;32(9):915–25.

28. Zhao W, He X, Hoadley KA, Parker JS, Hayes DN, Perou CM. Comparison of RNA-Seq by poly (A) capture, ribosomal RNA depletion, and DNA microarray for expression profiling. BMC Genomics. 2014;15(1):419.

29. Cieslik M, Chugh R, Wu Y-M, Wu M, Brennan C, Lonigro R, et al. The use of exome capture RNA-seq for highly degraded RNA with application to clinical cancer sequencing. Genome Res. 2015/08/07. 2015 Sep;25(9):1372–81.

30. Ryan MM, Guévremont D, Mockett BG, Abraham WC, Williams JM. Circulating Plasma microRNAs are Altered with Amyloidosis in a Mouse Model of Alzheimer’s Disease. J Alzheimers Dis. 2018;66(2):835–52.

31. Andrews S. FastQC: A Quality Control Tool for High Throughput Sequence Data. 2015 Jun;

32. Schubert M, Lindgreen S, Orlando L. AdapterRemoval v2: rapid adapter trimming, identification, and read merging. BMC Res Notes. 2016 Dec;9(1):88.

33. Bolger AM, Lohse M, Usadel B. Trimmomatic: A flexible trimmer for Illumina sequence data. Bioinformatics. 2014 Aug;30(15):2114–20.

34. Dobin A, Davis CA, Schlesinger F, Drenkow J, Zaleski C, Jha S, et al. STAR: Ultrafast universal RNA-seq aligner. Bioinformatics. 2013 Jan;29(1):15–21.

35. Friedländer MR, MacKowiak SD, Li N, Chen W, Rajewsky N. MiRDeep2 accurately identifies known and hundreds of novel microRNA genes in seven animal clades. Nucleic Acids Res. 2012 Jan;40(1):37–52.

36. Sai lakshmi S, Agrawal S. piRNABank: A web resource on classified and clustered Piwi-interacting RNAs. ucleic Acids Res. 2008 Jan;36(SUPPL. 1):D173.

37. Robinson MD, McCarthy DJ, Smyth GK. edgeR: a Bioconductor package for differential expression analysis of digital gene expression data. Bioinformatics. 2010 Jan;26(1):139–40.

38. Liao Y, Smyth GK, Shi W. The R package Rsubread is easier, faster, cheaper and better for alignment and quantification of RNA sequencing reads. Nucleic Acids Res. 2019 May;47(8):e47–e47.

39. Morgan M, Pagès H, Obenchain V, Hayden N. Rsamtools: Binary alignment (BAM), FASTA, variant call (BCF), and tabix file import. R package version 2.4.0. 2020.

40. Wickham H. stringr: Sample, Consistent Wrappers for Common String Operations. 2019.

41. Wickham H. ggplot2: Elegant Graphics for Data Analysis. New York: Springer-Verlag; 2016.

42. Bengtsson H. matrixStats: Functions that Apply to Rows and Columns of Matrices (and to Vectors). 2020.

43. Kolde R. pheatmap: Pretty Heatmaps. 2019.

44. Wickham H, Averick M, Bryan J, Chang W, McGowan L, François R, et al. Welcome to the Tidyverse. J Open Source Softw. 2019 Nov;4(43):1686.

45. Edgar R, Domrachev M, Lash AE. Gene Expression Omnibus: NCBI gene expression and hybridization array data repository. Nucleic Acids Res [Internet]. 2002 Jan 1;30(1):207–10. Available from: https://pubmed.ncbi.nlm.nih.gov/11752295

46. McDermaid A, Chen X, Zhang Y, Wang C, Gu S, Xie J, et al. A New Machine Learning-Based Framework for Mapping Uncertainty Analysis in RNA-Seq Read Alignment and Gene Expression Estimation. Vol. 9, Frontiers in Genetics. 2018. p. 313.

47. Dharshini SAP, Taguchi Y-H, Gromiha MM. Identifying suitable tools for variant detection and differential gene expression using RNA-seq data. Genomics. 2020;112(3):2166–72.

48. Gregory TR. The C-value enigma in plants and animals: a review of parallels and an appeal for partnership. Ann Bot. 2005 Jan;95(1):133–46.

49. Palazzo AF, Lee ES. Non-coding RNA: what is functional and what is junk? Vol. 6, Frontiers in Genetics. 2015. p. 2.

50. O’Reilly D, Dienstbier M, Cowley SA, Vazquez P, Drozdz M, Taylor S, et al. Differentially expressed, variant U1 snRNAs regulate gene expression in human cells. Genome Res. 2012/10/15. 2013 Feb;23(2):281–91.

51. Dvinge H, Guenthoer J, Porter PL, Bradley RK. RNA components of the spliceosome regulate tissue- and cancer-specific alternative splicing. Genome Res. 2019 Oct;29(10):1591–604.

52. Shagin DA, Lukyanov KA, Vagner LL, Matz M V. Regulation of average length of complex PCR product. Nucleic Acids Res. 1999 Sep;27(18):e23-i–e23-iii.

53. Boivin V, Deschamps-Francoeur G, Couture S, Nottingham RM, Bouchard-Bourelle P, Lambowitz AM, et al. Simultaneous sequencing of coding and noncoding RNA reveals a human transcriptome dominated by a small number of highly expressed noncoding genes. RNA. 2018/04/27. 2018 Jul;24(7):950–65.

54. Nottingham RM, Wu DC, Qin Y, Yao J, Hunicke-Smith S, Lambowitz AM. RNA-seq of human reference RNA samples using a thermostable group II intron reverse transcriptase. RNA. 2016 Apr;22(4):597–613.

55. Qin Y, Yao J, Wu DC, Nottingham RM, Mohr S, Hunicke-Smith S, et al. High-throughput sequencing of human plasma RNA by using thermostable group II intron reverse transcriptases. RNA. 2016 Jan;22(1):111–28.

56. Xu H, Yao J, Wu DC, Lambowitz AM. Improved TGIRT-seq methods for comprehensive transcriptome profiling with decreased adapter dimer formation and bias correction. Sci Rep. 2019 May;9(1):7953.

57. Yao J, Wu DC, Nottingham RM, Lambowitz AM. Identification of protein-protected mRNA fragments and structured excised intron RNAs in human plasma by TGIRT-seq peak calling. Elife. 2020 Sep;9.

